# Predicting disease-overarching therapeutic approaches for Congenital Disorders of Glycosylation using multi-OMICS

**DOI:** 10.1101/2025.07.07.663468

**Authors:** I.J.J. Muffels, R. Budhraja, R. Shah, S. Radenkovic, E. Morava, T Kozicz

**Affiliations:** Department of Genetics and Genomics, Icahn school of Medicine at Mount Sinai, New York, NY, USA; Department of Laboratory Medicine and Pathology, Mayo Clinic, Rochester, Minnesota, United States; Zydus research center, Zydus Lifesciences Limited, Ahmedabad, Gujarat, India; Department of Metabolic Diagnostics, University Medical Center Utrecht, Utrecht, The Netherlands; Department of Biophysics, University of Pecs Medical School, Pecs, Hungary; Department of Anatomy, University of Pecs Medical School, Pecs, Hungary

## Abstract

**Background:** Congenital Disorders of Glycosylation (CDG) are a rapidly expanding group of inherited metabolic diseases caused by defects in glycosylation. Although over 190 genetic defects have been identified, effective treatments remain available for only a few. We hypothesized that integrative analysis of multi-omics datasets from individuals with various CDG could uncover common molecular signatures and highlight shared therapeutic targets.

**Methods:** We compiled all publicly available RNA sequencing, proteomics and glycoproteomics datasets from patients with PMM2-CDG, ALG1-CDG, SRD5A3-CDG, NGLY1-CDDG, ALG13-CDG and PGM1-CDG, spanning different tissues, including induced cardiomyocytes, human cortical organoids, fibroblasts, and lymphoblasts. Differential expression and glycosylation analyses were performed, followed by Gene Set Enrichment Analysis (GSEA) to identify commonly dysregulated pathways. We then applied the EMUDRA drug prediction algorithm to prioritize candidate compounds capable of reversing these shared molecular signatures.

**Results:** We identified four glycoproteins with consistent differential glycosylation across all eight glycoproteomics datasets. Six glycosylation sites and glycan structures were recurrently altered across CDG and showed partial correction with treatment. Pathway analysis revealed shared disruptions in autophagy, vesicle trafficking, and mitochondrial function. EMUDRA predicted several repurposable drug classes, including muscle relaxants, antioxidants, beta-adrenergic agonists, antibiotics, and NSAIDs, that could reverse key pathway abnormalities, particularly those involving autophagy and N-glycosylation.

**Conclusion:** Most dysregulated pathways were shared across CDG, suggesting the potential for common therapeutic strategies. Several candidate drugs targeting these shared abnormalities emerged from integrative analysis and warrant validation in future in vitro studies.

## Introduction

Congenital disorders of glycosylation (CDG) result from impaired glycosylation and are categorized according to their primarily affected glycosylation type: N-glycosylation, O-glycosylation, combined glycosylation defects, glycolipid synthesis, or other related pathway abnormalities (Lefeber et al., 2022) CDG have been associated with pathogenic variants in over 190 different genes.(Ng et al., 2024) Protein glycosylation is highly abundant, with over 50% of proteins being glycosylated.(Apweiler et al., 1999) Glycosylated proteins are involved in a plethora of cellular functions, and contribute to proper protein folding and function, extracellular matrix structure, energy metabolism and cellular signaling.(Gagneux et al., 2022) The clinical phenotype of CDG is highly heterogenous. This might be due to the multitude of dysregulated pathways observed in these diseases. (Ligezka et al., 2023; Zdrazilova et al., 2023)

Currently available treatment options for CDG are mostly symptomatic. Dietary supplementation of monosaccharides has been used for many years, although it is only effective for a subset of CDG. (Verheijen et al., 2020) Recently, large-scale drug screenings of FDA-approved drugs have come into play, providing a novel approach to discover therapies for CDG. (Dalton et al., 2024; Iyer et al., 2019; Ligezka et al., 2021; Radenkovic et al., 2023) Drug screening is usually performed by introducing a specific genetic defect in simple disease models (yeast, worm or fly), and verifying which compounds increase overall growth or survival. However, as these models are tailored to a specific genetic defect or variant, the results might not be translatable to other patients or other CDG. Developing novel treatments by modeling each of the 190 known CDG-associated genes individually is a time-intensive endeavor that could take years. Moreover, most candidate therapies require validation in patient-derived cells or human model systems before advancing to clinical use.(Muffels et al., 2025) Generating distinct human models for every CDG is both technically challenging and cost-prohibitive. Identifying a single drug—or class of drugs—that targets shared disease mechanisms across multiple CDG would significantly accelerate therapeutic development by enabling efficient drug repurposing.

Over the past decade, OMICS approaches have enabled the comprehensive analysis of an entire class of biomolecules, such as proteins, RNA, or glycan species. Thus far, several CDG have been characterized using these OMICS approaches.(Budhraja, Joshi, et al., 2024; de Haas et al., 2022; Garapati et al., 2024; Ligezka et al., 2023; Radenkovic et al., 2024; Wessels et al., 2024) OMICS datasets offer valuable insights into shared pathways across different genetic defects, highlighting potential targets for therapeutic development. Additionally, OMICS data can be leveraged to predict effective therapies using existing libraries that evaluate the effects of various FDA-approved drugs in cell lines.(Lamb et al., 2006; Zhou et al., 2018) Thus, comparing OMICS data across CDG offers dual advantages: it enhances understanding of shared phenotypes among different CDG and facilitates the prediction of repurposed drugs that could serve as novel therapeutic options.

In this study, we integrated RNA sequencing, proteomics, and glycoproteomics datasets to identify shared and specific molecular pathways across six CDG: PMM2-CDG, NGLY1-CDDG, SRD5A3-CDG, ALG1-CDG, ALG13-CDG, and PGM1-CDG, using three distinct human disease model systems.

Following pathway analysis, we applied the EMUDRA drug prediction algorithm to prioritize potential therapeutic candidates. This integrative OMICS-based approach highlights the power of systems-level analysis to uncover convergent disease mechanisms and identify broadly applicable treatment strategies across diverse CDG.

## Methods

Proteomics, RNA-sequencing and glycoproteomics data were extracted from literature.(Budhraja et al., 2023; Budhraja, Joshi, et al., 2024; Garapati et al., 2024; Ligezka et al., 2023; Parrado et al., 2022; Radenkovic et al., 2024, 2025) ALG13-CDG data was extracted from the PRIDE repository (PXD05647).

### Differentially Expressed (Glyco)proteins

To compare differentially expressed (glyco-)proteins between different diseases, the data was filtered for entries that were significantly different between healthy controls and individuals with CDG. (Glyco)proteins with unadjusted p < 0.1 (not corrected for multiple testing) were considered candidates in each dataset; consistency across datasets was then assessed as an additional criterion to reduce the likelihood of false positives. All differentially expressed glycoproteins (regardless of the glycan species attached to it) were compared between diseases to create **Figure 1A**. CDG patient-derived fibroblasts were compared before and after treatment with either Epalrestat or GLM101 to create

**Figure 1.**
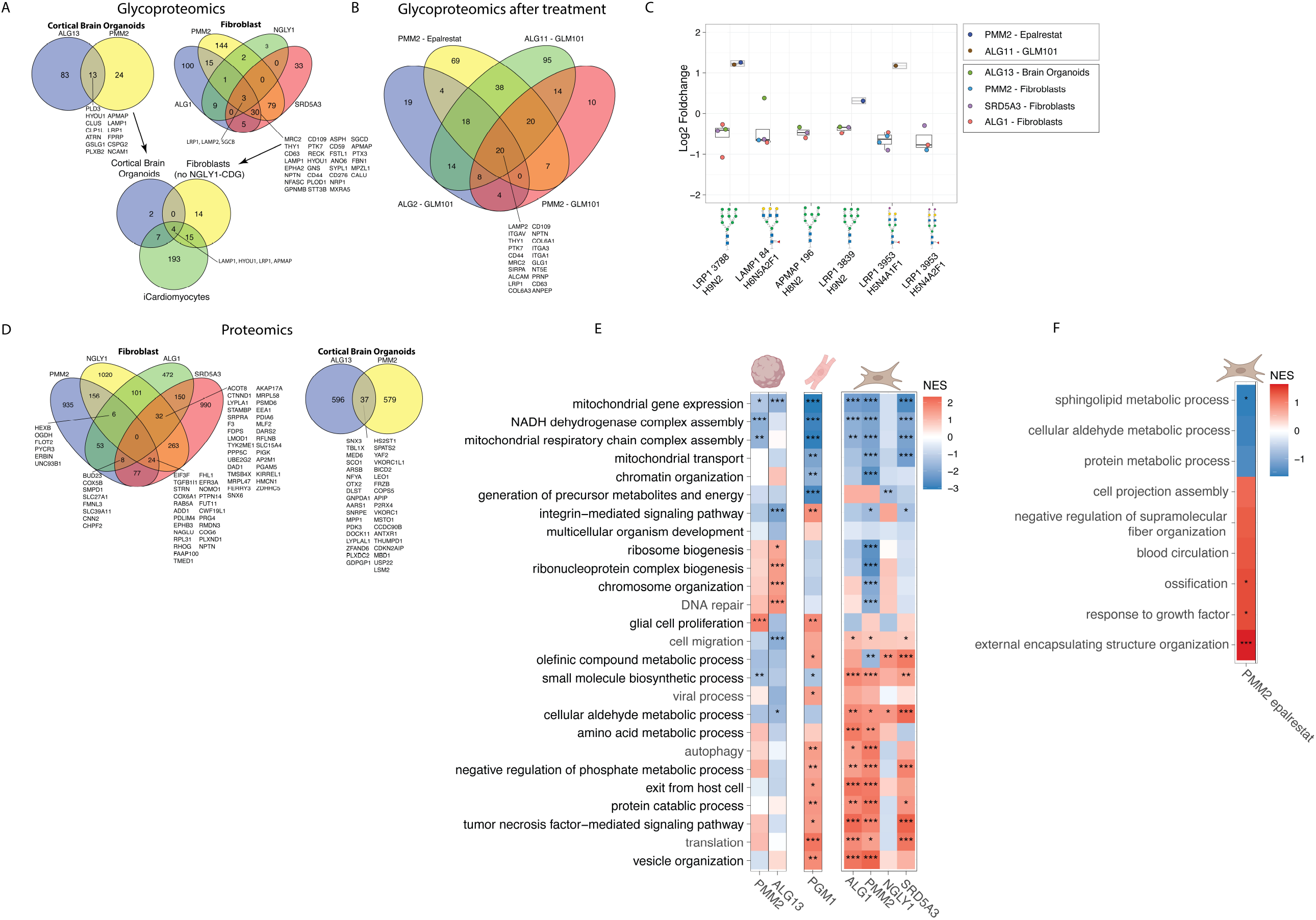
Overlapping pathways among different CDG. (A) Showing overlapping glycoproteins with significantly different expression in patients compared to healthy controls (p-value <0.1) across CDG and tissues. (B) Showing overlapping glycoproteins with significantly different expression in untreated patient-derived cells compared to treated patient-derived cells (p-value <0.1). (C) Displaying the log2 foldchange values of four specific glycoproteins that were differentially glycosylated in at least three or more glycoproteomics datasets. All of these specific glycoproteins were downregulated in CDG patients but could be upregulated during treatment. (D) Showing overlapping differentially expressed proteins in patients compared to healthy controls (p-value <0.1) across CDG and tissues. (E) Heat map showing the Gene Set Enrichment Analysis results for the different proteomics datasets. NES refers to Normalized Enrichment Score. The tissues in which the proteomics where performed are shown graphically above the heat map. p<0.05*, p<0.01**, p<0.001***, p<0.0001****. (F) Showing Gene Set Enrichment Analysis results for the proteomics results of PMM2-CDG fibroblasts treated with Epalrestat.

**Figure 1B. Figure 1C** was created by focusing on the four differentially glycosylated proteins that were shared across diseases from **Figure 1A**, i.e. sites that were differentially glycosylated with the same species, across different CDG are highlighted in **Figure 1C**. For **Figure 1E**, all differentially expressed proteins shared between diseases are shown.

### Gene Set Enrichment Analysis

To create **Figure 1E**, Gene Set Enrichment Analysis was performed. First, the different proteomics datasets were ranked based on the log2 foldchange of individuals with CDG compared to controls. Subsequently, GSEA was performed using the ClusterProfiler Package in R, using FDR to correct for multiple testing. Redundant ontologies were merged based on the GO ontology hierarchy, where the pathways further up the tree were taken as an overarching category **(Supplementary Table 2)**. For GO ontologies belonging to the same category, a mean p-value and Normalized Enrichment Score (NES) were calculated, which are shown in **Figure 1E** In addition, we focused solely on pathways with similar p-values between CDG, which were filtered by taking all ontologies with differences in p-value (standard deviation) <0.2. The individual p-values and NES can be found in **Supplementary Table 2**. The stars are based on the raw p-values: p<0.05*, p<0.01**, p<0.001***, p<0.0001****.

### Ensemble of Multiple Drug Repositioning Approach (EMUDRA)

To create **Figure 2**, EMUDRA was performed based on the differentially expressed proteins or genes of the different OMICS datasets. Cutoff values for EMDURA were unadjusted p<0.05 and log2foldchange >0.25, except for ALG13-CDG cortical brain organoid data, where the cutoff value was log2foldchange >0.2. The compounds with the lowest Normalized Prediction Scores, thereby having stronger ability to reverse the transcriptomic phenotype to normal values, were used for downstream analysis. Biochemical Drug classes were attributed to these top-20 compounds through searching the ChebI Ontology.(Hastings et al., 2016) Functional classes and disease indications were attributed to the compounds by searching the Drugbank database. (Wishart et al., 2006)

**Figure 2.**
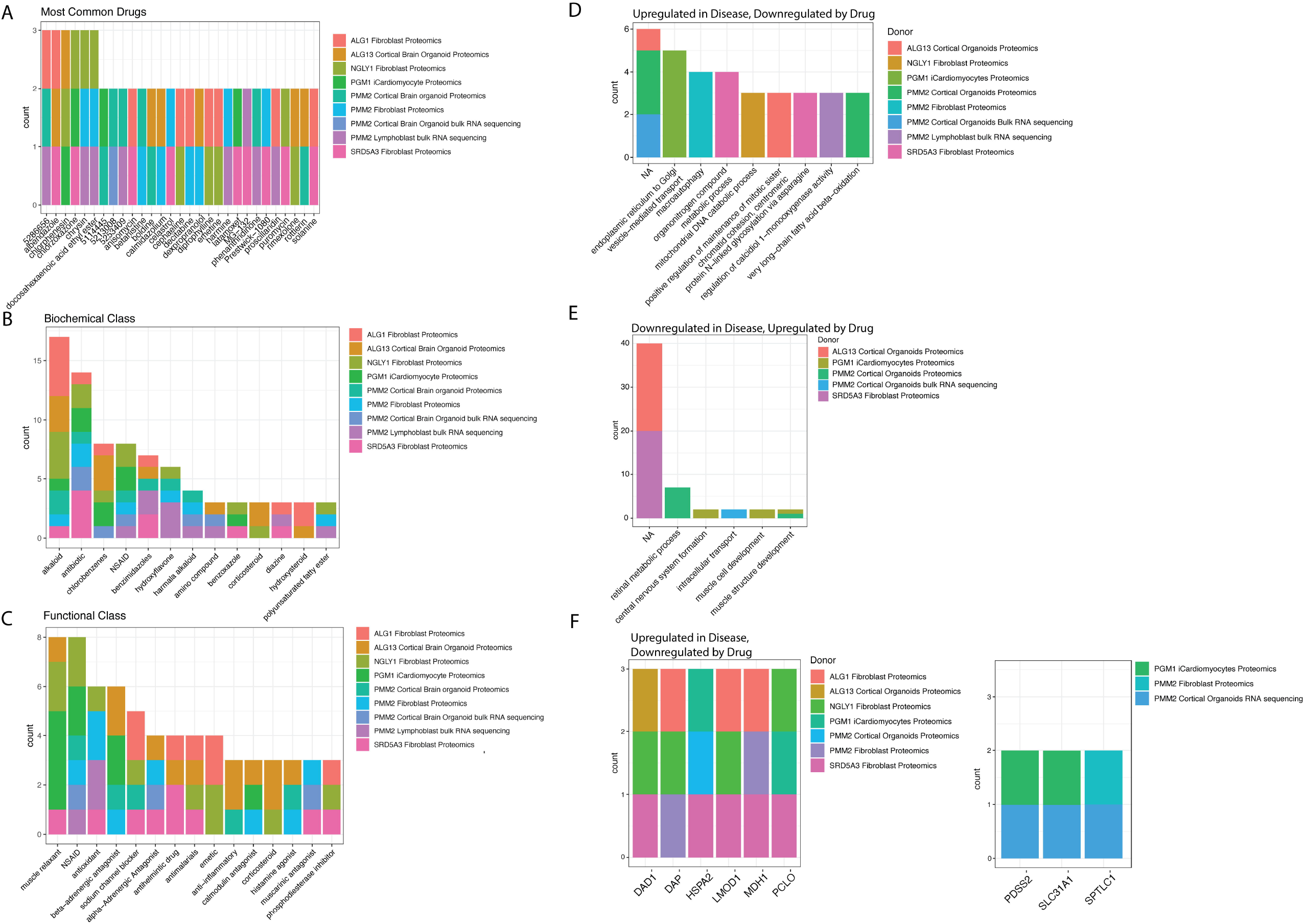
Showing the results of the Ensemble of Drug Repurposing Approaches (EMUDRA) (A) For each CDG, we focused on the top-20 drugs identified with EMUDRA. We counted the frequency of each drug within these datasets, shown on the Y-axis. The colors refer to the individual CDG and tissues. (B) Displaying the biochemical classes to which the top-20 drugs identified with EMUDRA belonged to. Biochemical classes were identified through the CheBI ontology. (C) Displaying the functional classes to which the top-20 drugs identified with EMUDRA belonged to. Functional classes were identified using the Drugbank database. (D) Showing the pathways that could be restored by the top-20 drugs identified with EMUDRA, which were downregulated in CDG patients. (E) Showing the pathways that were downregulated by the top-20 drugs identified with EMUDRA, which were upregulated in CDG patients. (F) Showing the genes that could be significantly upregulated or downregulated by the top-20 drugs identified by EMUDRA. Each gene was only counted once per dataset.

## Results

We extracted OMICS data from human cortical organoids (ALG13-CDG, PMM2-CDG), iCardiomyocytes (PGM1), fibroblasts (PMM2-CDG, SRD5A3-CDG, ALG1-CDG) and lymphoblasts (PMM2-CDG) **(Table 1)**. We also included proteomics data of a deglycosylation disease, derived from fibroblasts of individuals with NGLY1-Congenital Disorder of Deglycosylation (CDDG). The majority of these datasets were proteomics-based. However, for PMM2-CDG human cortical organoids and PMM2-CDG lymphoblasts, we also had bulk RNA sequencing data available. The number of patients differed for each dataset, ranging from two to five patients **(Table 1)**. In addition, we also included glycoproteomics and proteomics data of patient-derived fibroblasts before and after different treatments that are currently trialed in CDG patients (Epalrestat (NCT04925960) and GLM101 (TRIAL 2024-513119-29-00)).

**Table 1:**
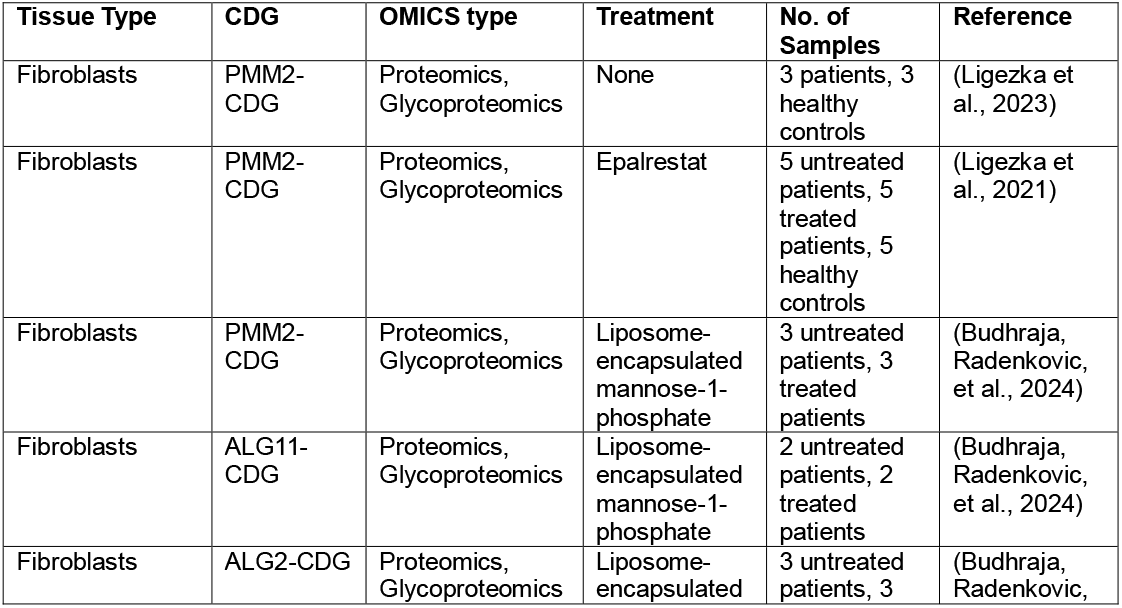
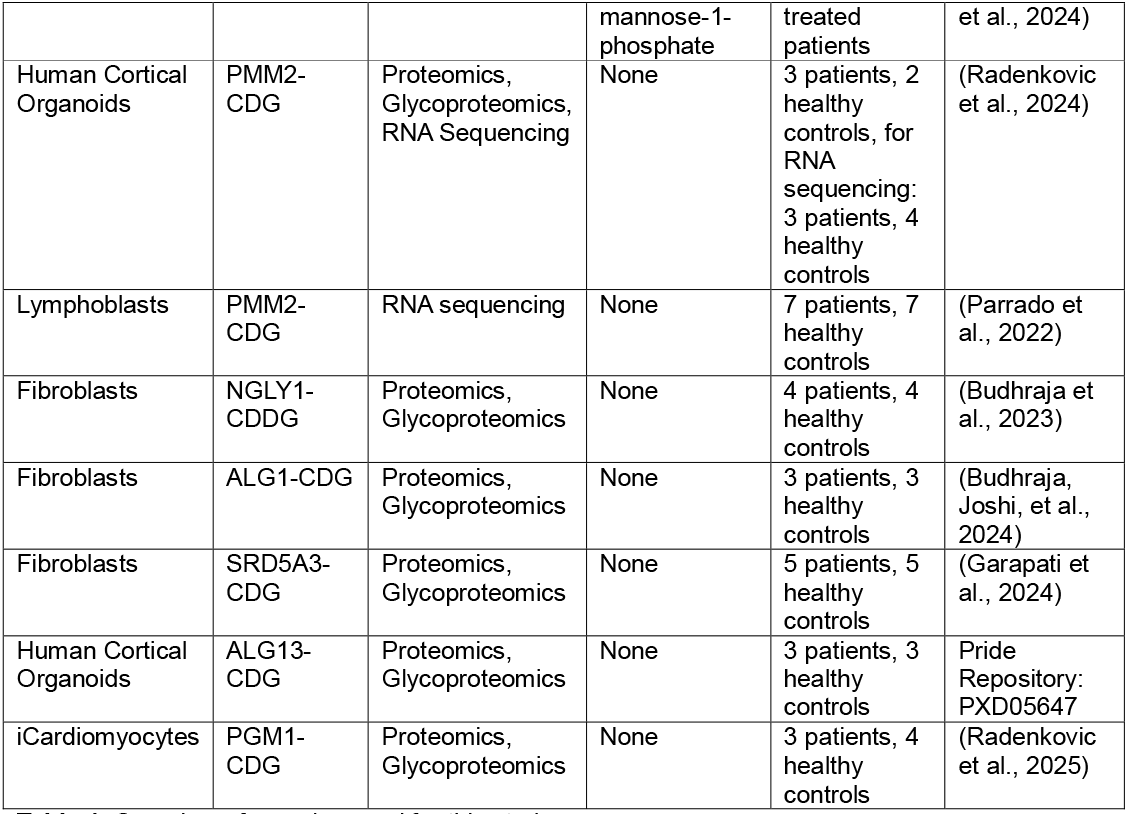
Overview of samples used for this study.

### Glycosylation of LRP1, LAMP1, HYOU1 and APMAP is aberrant across all CDG and is responsive to treatment

First, we assessed the overlap between genetic defects and tissues regarding differentially expressed glycoproteins **(Figure 1A)**. Within human cortical organoids of ALG13-CDG and PMM2-CDG, 13 proteins showed significantly different glycosylation and were shared, while only three glycoproteins (LAMP2, LRP1, SGCB) were shared between PMM2-CDG, ALG1-CDG, SRD5A3-CDG and NGLY1-CDDG fibroblasts. Since pathogenic variants in *NGLY1* lead to a deglycosylation disease, we also assessed overlapping glycan species when we removed the NGLY-CDDG dataset from the comparison, which resulted in 30 overlapping differentially glycosylated proteins, including THY1, STT3B and LAMP1, in fibroblasts.

When comparing datasets across different tissue types—human cortical organoids, iCardiomyocytes, and fibroblasts (excluding the NGLY1-CDDG dataset) — we identified four glycoproteins (LAMP1, HYOU1, LRP1, and APMAP) that consistently exhibited aberrant glycosylation (**Figure 1A**). Among these, only LRP1 also showed differential glycosylation in the NGLY1-CDDG dataset. Notably, the specific glycan structures and glycosylation site positions on these proteins varied across tissues. Only six glycan species were consistently attached to the same site across multiple datasets, indicating limited overlap at the level of site-specific glycan composition between CDG (**Figure 1C**).

We also assessed which glycoproteins and glycan species changed when fibroblasts were treated with either Epalrestat or GLM101 **(Figure 1B, C)**. Both treatments are currently trialed clinically in PMM2-CDG patients and showed beneficial therapeutic effects *in-vitro*. (Budhraja, Radenkovic, et al., 2024; Glycomine, 2022; Ligezka et al., 2021) Intriguingly, several of these shared glycan species changed significantly during these treatments, indicating that these glycan species might serve as relevant biomarkers for multiple CDG.

Taken together, we identified several proteins that are almost always aberrantly glycosylated across CDG. In addition, we identified several specific glycan biomarkers that are responsive to treatment and may serve as CDG biomarkers.

### Differentially expressed proteins across CDG

Next, we focused on proteomics data, by assessing which proteins were differentially expressed in the different CDG and the pathways they were involved in. While the human cortical organoids of ALG13-CDG and PMM2-CDG shared 37 differentially expressed proteins, the four fibroblast datasets shared none. We also assessed the overlap between SRD5A3-CDG, PMM2-CDG and ALG1-CDG – excluding NGLY1-CDDG - revealing eight overlapping differentially expressed proteins **(Figure 1D)**.

To assess which pathways were significantly enriched in the datasets, we performed Gene Set Enrichment Analysis (GSEA). Interestingly, we found that mitochondrial translation, oxidative phosphorylation and the respiratory electron chain proteins were downregulated in all CDG, among all the different tissues assessed (cardiomyocytes, human cortical organoids, fibroblasts) **(Figure 1E)**. In contrast, we found autophagy was upregulated in all CDG, except for ALG13-CDG and NGLY1-CDDG. Collagen Fibril Organization and Extracellular Matrix Remodeling were significantly downregulated in CDG-derived human cortical organoids, but not in other tissues. We also assessed the effects of Epalrestat treatment on proteomics results. Epalrestat significantly increased extracellular matrix organization (which was grouped under external encapsulating structure organization) **(Figure 1F)**.

Overall, the integration of proteomics and pathway analyses revealed both CDG-specific and tissue-specific pathophysiological mechanisms. While mitochondrial dysfunction and autophagy emerged as a common feature across CDG and tissues, extracellular matrix remodeling varied depending on the specific CDG and the tissue analyzed.

### Prediction of novel treatment targets based on multi-OMICS datasets

Next, we employed EMUDRA to predict novel treatment targets based on the different OMICS datasets that were included in this study. An overview of the results can be found in **Supplementary Table 1**. We performed EMUDRA on either proteomics and/or transcriptomics for different tissues and CDG. Apart from the datasets shown in **Figure 1**, we also assessed transcriptomics data derived from PMM2-CDG lymphoblasts and human cortical organoids.

Several compounds were able to partially normalize gene expression profiles across at least three independent datasets. Notably, compound 5286656 emerged as a top candidate in PMM2-CDG patient-derived lymphoblasts, human cortical organoids, and ALG1-CDG patient-derived fibroblasts **(Figure 2A)**. Although its exact mechanism of action remains unknown, 5286656 modulated key disease-relevant pathways in a context-specific manner. For example, it downregulated the upregulated “protein deglycosylation involved in glycoprotein catabolic process” pathway in PMM2-deficient lymphoblasts, and restored “vitamin A metabolism,” which was downregulated in PMM2-CDG human cortical organoids.

Chlorphenesin was predicted for NGLY1-CDDG, ALG13-CDG and PGM1-CDG. Although its exact mechanism of action is not known, it might have dampening effects on certain aspects of CNS signaling.(Aihara et al., 1980) Chrysin is a compound extracted from passion fruit and was predicted to help NGLY1-CDDG and PMM2-CDG, potentially due to its antioxidant and anti-inflammatory effects. (Del Fabbro et al., 2025) Docosahexaenoic acid (DHA) ethyl ester, predicted to help NGLY1-CDDG and PMM2-CDG, is an esterified version of the omega-3 fatty acid DHA. DHA is incorporated into cellular membranes, for example mitochondrial membranes, where it aids mitochondrial function. (Stillwell et al., 1997)

To better understand how the predicted drugs might improve cellular phenotypes in CDG, we grouped the candidate compounds by drug class **(Figure 2B/C)**. Alkaloids—naturally occurring compounds containing at least one nitrogen atom—emerged as top candidates across all CDG, likely reflecting the abundance of alkaloids in the FDA-approved drug library. Additionally, antibiotics, NSAIDs, antioxidants, and chlorobenzenes were identified as promising candidates in four of the five CDG assessed, suggesting potential for broader therapeutic relevance across the CDG-spectrum.

EMUDRA not only provides promising drug targets for CDG, but it also indicates which pathways are restored under each treatment **(Figure 2D)**. During treatment, several pathways could be normalized, such as restoration of autophagy for PMM2-CDG (fibroblasts), and vesicle mediated transport for PGM1-CDG (iCardiomyocytes) (**Supplementary Table 1)**. Intriguingly, similar drugs normalized different pathways, depending on the CDG involved **(Supplementary Table 1)**. EMUDRA not only predicts pathways that normalize under treatment, it also predicts which genes can be normalized **(Figure 2E)**. In various CDG, EMUDRA identified drugs that normalized the expression of the Oligosaccharyltransferase (OST) complex subunit DAD1 and heatshock protein HSPA2 **(Figure 2E)**. EMUDRA also identified drugs that could upregulate the expression of PDSS2, a protein that uses farnesyl diphosphate (FPP) and isopentenyl diphosphate (IPP) to produce the prenyl sidechain of coenzyme Q, which is essential for complex II functionality (López et al., 2006).

Together, our findings demonstrate that EMUDRA can predict potential treatment targets across multiple CDG, with several drug classes emerging as promising candidates for normalizing disease-associated gene expression profiles. Notably, antibiotics, NSAIDs, antioxidants and chlorobenzenes were identified as potential therapies, acting through diverse mechanisms such as reducing autophagy for PMM2-CDG.

## Discussion

In this study, we integrated multi-omics data from diverse genetic etiologies of CDG to identify shared pathogenic mechanisms and potential therapeutic strategies. Our analyses revealed that mitochondrial dysfunction, extracellular matrix remodeling, and dysregulated autophagy are common features across multiple CDG types. Glycoproteomics data further showed consistent aberrant glycosylation of LRP1, LAMP1, HYOU1, and APMAP across different tissues, with some glycan species and modification sites conserved across CDG. Notably, these glycosylation patterns partially normalized following treatment, suggesting their potential as biomarkers for diagnostics and therapeutic response. Drug prediction using EMUDRA identified several compounds with efficacy across multiple CDG, including muscle relaxants, antioxidants, beta-adrenergic agonists, antibiotics, and NSAIDs. These findings offer a promising foundation for the development of broadly applicable therapies for multiple CDG.

Several of the identified drugs align with previous attempts at treating CDG, or known data on underlying pathophysiology. For instance, antioxidants were identified as a key treatment target by EMUDRA, correlating with increased levels of oxidative stress found in CDG, including ATP6V1A-CDG, PMM2-CDG, ALG8-CDG, RFT1-CDG, and SLC10A7-CDG.(Ondruskova et al., 2020; Sun et al., 2025) Antioxidants were also identified as potential treatments in PMM2-CDG worm- and yeast models.(Iyer et al., 2019; Lao et al., 2019) The identification of NSAIDs was intriguing, as they inhibit the COX2 enzyme, whose expression was found to be upregulated in PMM2-CDG patient-derived fibroblasts.(Gallego et al., 2024) NSAIDs were also identified as targets in an FDA-approved drug screening in a fly model of MAN1B1-CDG, and ibuprofen treatment or COX2 inhibition led to a significant reduction of seizures.(Chow et al., 2024) Chrysin, a compound predicted to improve PMM2-CDG and NGLY1-CDDG, has similarly been shown to reduce COX2 activity.(Del Fabbro et al., 2025) Interestingly, Hydroxyflavones were predicted as top hits to treat PMM2-CDG, and came up in all four PMM2-CDG datasets. In PMM2 mutant worms, hydroxyflavones were similarly identified as top hits in large-scale FDA drug screenings (Iyer et al., 2019). Hydroxyflavones are known for their antioxidant function and have been shown to increase PMM2 activity in fibroblasts.(Iyer et al., 2019) Harmala alkaloids were also predicted as targets for PMM2-CDG datasets specifically. Harmala alkaloids inhibit monoamino oxidase A, which breaks down dopamine, norepinefrine and serotonine. Intriguingly, levodopa and ethylnorepinephrine came up as top hits to treat PMM2 worms as well.(Iyer et al., 2019) The effect of hydroxyflavones and harmala alkaloids might be quite specific for PMM2, as these compounds were not identified in any other CDG in this manuscript. Antibiotics, muscle relaxants, beta-adrenergic antagonist and alfa-adrenergic agonists came up frequently for various types of CDG, and could normalize various pathways, including the actin cytoskeleton and vesicle trafficking. Of note, propranolol, a beta-adrenergic antagonist, has been found to induce accumulation of intracellular glycoproteins in muscle cells, potentially due to altered endosomal trafficking, which may offer insight into its mechanism of action in the CDG context.(Han et al., 2016)

Our findings support the feasibility of basket trials for CDG, given the significant overlap in dysregulated pathways across multiple CDG. Basket trials—designed to evaluate a single therapy in patients with distinct genetic mutations but shared pathophysiology—offer a powerful strategy to accelerate drug development for rare disorders like CDG (Muffels et al., 2025) However, without functional validation, the efficacy and safety of the predicted compounds in this manuscrupt remain uncertain. Future studies should point out whether the candidate drugs are effective in relevant disease models of CDG patients. Looking ahead, the integration of multi-omics-driven drug prediction into rare disease research offers a promising framework. By harnessing large-scale datasets, researchers can more precisely identify candidate drugs with the highest likelihood of success—and flag those with potential adverse effects based on the predicted pathways where they interfere in. This strategy not only streamlines the drug discovery process but also deepens our understanding of the molecular pathways driving disease, guiding the development of more targeted therapies for CDG and other metabolic disorders.

## Limitations

EMUDRA relies on transcriptomic Connectivity Map (CMAP) data derived from cancer cell lines, which may not fully capture the complexity of tissues relevant to CDG pathophysiology. In addition, the application of proteomics data on a transcriptomic network is not ideal because protein abundance and activity are governed by post⍰transcriptional regulation, postliltranslational modifications, and protein–protein interactions that are not reflected in mRNA expression maps, leading to potential mismatches between observed protein changes and the underlying transcriptlilbased connectivity.

Additionally, the algorithm is designed to restore as many gene expression changes as possible toward baseline, often yielding broad and non-specific pathway predictions. Importantly, EMUDRA does not distinguish between pathogenic and compensatory gene expression changes. For example, in SRD5A3-CDG fibroblasts, upregulation of OST complex subunits—likely a compensatory response—led to the identification of drugs predicted to downregulate N-glycosylation, a strategy that could be ineffective or even harmful in the context of CDG. Therefore, the therapeutic candidates predicted by EMUDRA must be carefully evaluated for context-specific effects and potential adverse outcomes before advancing to preclinical testing.

## Supporting information

Overview of EMDURA results for all datasets included in this manuscript.

Overview of child and parent GO ontologies used for Figure 1E.

## Acknowledgements

This study was supported by 1U54NS115198-01 from the National Institute of Neurological Diseases and Stroke (NINDS), the National Center for Advancing Translational Sciences (NCATS), the National Institute of Child Health and Human Development (NICHD), and the Rare Disorders Consortium Disease Network (RDCRN).

## References

Aihara, H., Kurachi, M., Nakane, S., Sasajima, M., & Ohzeki, M. (1980). THE ACTION OF CHLORPHENESIN CARBAMATE ON THE FROG SPINAL CORD. Japanese Journal of Pharmacology, 30(1), 29–36. 10.1254/jjp.30.29

Apweiler, R., Hermjakob, H., & Sharon, N. (1999). On the frequency of protein glycosylation, as deduced from analysis of the SWISS-PROT database. Biochimica et Biophysica Acta (BBA) - General Subjects, 1473(1), 4–8. 10.1016/S0304-4165(99)00165-8

Budhraja, R., Joshi, N., Radenkovic, S., Kozicz, T., Morava, E., & Pandey, A. (2024). Dysregulated proteome and N-glycoproteome in ALG1-deficient fibroblasts. PROTEOMICS, 24(15). 10.1002/pmic.202400012

Budhraja, R., Radenkovic, S., Jain, A., Muffels, I. J. J., Ismaili, M. H. A., Kozicz, T., Pandey, A., & Morava, E. (2024). Liposome-encapsulated mannose-1-phosphate therapy improves global N-glycosylation in different congenital disorders of glycosylation. Molecular Genetics and Metabolism, 142(2), 108487. 10.1016/j.ymgme.2024.108487

Budhraja, R., Saraswat, M., De Graef, D., Ranatunga, W., Ramarajan, M. G., Mousa, J., Kozicz, T., Pandey, A., & Morava, E. (2023). N-glycoproteomics reveals distinct glycosylation alterations in <scp>NGLY1</scp> -deficient patient-derived dermal fibroblasts. Journal of Inherited Metabolic Disease, 46(1), 76–91. 10.1002/jimd.12557

Chow, C., Beebe, L., Hope, K. A., Coelho, E., Evans, H. D., Massey, C., Fast, C., & Perlstein, E. O. (2024). A drug repurposing screen identifies NSAIDs and COX1/2 enzyme inhibition as potential therapies for MAN1B1-CDG, a rare congenital disorder of glycosylation. (ASHG 2024 Annual Meeting, Ed.; pp. 168–169).

Dalton, H. M., Young, N. J., Berman, A. R., Evans, H. D., Peterson, S. J., Patterson, K. A., & Chow, C. Y. (2024). A drug repurposing screen reveals dopamine signaling as a critical pathway underlying potential therapeutics for the rare disease DPAGT1-CDG. PLOS Genetics, 20(10), e1011458. 10.1371/journal.pgen.1011458

de Haas, P., de Jonge, M. I., Koenen, H. J. P. M., Joosten, B., Janssen, M. C. H., de Boer, L., Hendriks, W. J. A. J., Lefeber, D. J., & Cambi, A. (2022). Evaluation of Cell Models to Study Monocyte Functions in PMM2 Congenital Disorders of Glycosylation. Frontiers in Immunology, 13. 10.3389/fimmu.2022.869031

Del Fabbro, L., Bortolotto, V. C., Ferreira, L. M., Sari, M. H. M., & Furian, A. F. (2025). Chrysin’s anti-inflammatory action in the central nervous system: A scoping review and an evidence-gap mapping of its mechanisms. European Journal of Pharmacology, 997, 177602. 10.1016/j.ejphar.2025.177602

Gagneux, P., Hennet, T., & Varki, A. (2022). Essentials of Glycobiology (4th Edition). Cold Spring Harbor Laboratory Press.

Gallego, D., Serrano, M., Cordoba-Caballero, J., Gámez, A., Seoane, P., Perkins, J. R., Ranea, J. A. G., & Pérez, B. (2024). Transcriptomic analysis identifies dysregulated pathways and therapeutic targets in PMM2-CDG. Biochimica et Biophysica Acta (BBA) - Molecular Basis of Disease, 1870(5), 167163. 10.1016/j.bbadis.2024.167163

Garapati, K., Ranatunga, W., Joshi, N., Budhraja, R., Sabu, S., Kantautas, K. A., Preston, G., Perlstein, E. O., Kozicz, T., Morava, E., & Pandey, A. (2024). N-glycoproteomic and proteomic alterations in SRD5A3-deficient fibroblasts. Glycobiology, 34(11). 10.1093/glycob/cwae076

Glycomine. (2022). Mannose-1-Phosphate Replacement Therapy: A Potential Treatment for PMM2-CDG.

Han, S.-O., Pope, R., Li, S., Kishnani, P. S., Steet, R., & Koeberl, D. D. (2016). A beta-blocker, propranolol, decreases the efficacy from enzyme replacement therapy in Pompe disease. Molecular Genetics and Metabolism, 117(2), 114–119. 10.1016/j.ymgme.2015.09.012

Hastings, J., Owen, G., Dekker, A., Ennis, M., Kale, N., Muthukrishnan, V., Turner, S., Swainston, N., Mendes, P., & Steinbeck, C. (2016). ChEBI in 2016: Improved services and an expanding collection of metabolites. Nucleic Acids Research, 44(D1), D1214–D1219. 10.1093/nar/gkv1031

Iyer, S., Sam, F. S., DiPrimio, N., Preston, G., Verheijen, J., Murthy, K., Parton, Z., Tsang, H., Lao, J., Morava, E., & Perlstein, E. O. (2019). Repurposing the aldose reductase inhibitor and diabetic neuropathy drug epalrestat for the congenital disorder of glycosylation PMM2-CDG. DMM Disease Models and Mechanisms, 12(11). http://www.embase.com/search/results?subaction=viewrecord&from=export&id=L2004178174%0A10.1242/dmm.040584

Lamb, J., Crawford, E. D., Peck, D., Modell, J. W., Blat, I. C., Wrobel, M. J., Lerner, J., Brunet, J.-P., Subramanian, A., Ross, K. N., Reich, M., Hieronymus, H., Wei, G., Armstrong, S. A., Haggarty, S. J., Clemons, P. A., Wei, R., Carr, S. A., Lander, E. S., & Golub, T. R. (2006). The Connectivity Map: Using Gene-Expression Signatures to Connect Small Molecules, Genes, and Disease. Science, 313(5795), 1929–1935. 10.1126/science.1132939

Lao, J. P., DiPrimio, N., Prangley, M., Sam, F. S., Mast, J. D., & Perlstein, E. O. (2019). Yeast Models of Phosphomannomutase 2 Deficiency, a Congenital Disorder of Glycosylation. G3 (Bethesda, Md.), 9(2), 413–423. 10.1534/g3.118.200934

Lefeber, D. J., Freeze, H. H., Steet, R., & Kinoshita, T. (2022). Congenital Disorders of Glycosylation. In Essentials of Glycobiology (4th ed.). Cold Spring Harbor Laboratory Press.

Ligezka, A. N., Budhraja, R., Nishiyama, Y., Fiesel, F. C., Preston, G., Edmondson, A., Ranatunga, W., Van Hove, J. L. K., Watzlawik, J. O., Springer, W., Pandey, A., Morava, E., & Kozicz, T. (2023). Interplay of Impaired Cellular Bioenergetics and Autophagy in PMM2-CDG. Genes, 14(8), 1585. 10.3390/genes14081585

Ligezka, A. N., Radenkovic, S., Saraswat, M., Garapati, K., Ranatunga, W., Krzysciak, W., Yanaihara, H., Preston, G., Brucker, W., McGovern, R. M., Reid, J. M., Cassiman, D., Muthusamy, K., Johnsen, C., Mercimek-Andrews, S., Larson, A., Lam, C., Edmondson, A. C., Ghesquière, B., … Morava, E. (2021). Sorbitol Is a Severity Biomarker for PMM2-CDG with Therapeutic Implications. Annals of Neurology, 90(6), 887–900. 10.1002/ana.26245

López, L. C., Schuelke, M., Quinzii, C. M., Kanki, T., Rodenburg, R. J. T., Naini, A., DiMauro, S., & Hirano, M. (2006). Leigh Syndrome with Nephropathy and CoQ10 Deficiency Due to decaprenyl diphosphate synthase subunit 2 (PDSS2) Mutations. The American Journal of Human Genetics, 79(6), 1125–1129. 10.1086/510023

Muffels, I. J. J., Kozicz, T., Perlstein, E. O., & Morava, E. (2025). The Therapeutic Future for Congenital Disorders of Glycosylation. Journal of Inherited Metabolic Disease, 48(2). 10.1002/jimd.70011

Ng, B. G., Freeze, H. H., Himmelreich, N., Blau, N., & Ferreira, C. R. (2024). Clinical and biochemical footprints of congenital disorders of glycosylation: Proposed nosology. Molecular Genetics and Metabolism, 142(1), 108476. 10.1016/j.ymgme.2024.108476

Ondruskova, N., Honzik, T., Vondrackova, A., Stranecky, V., Tesarova, M., Zeman, J., & Hansikova, H. (2020). Severe phenotype of <scp>ATP6AP1-CDG</scp> in two siblings with a novel mutation leading to a differential tissue-specific <scp>ATP6AP1</scp> protein pattern, cellular oxidative stress and hepatic copper accumulation. Journal of Inherited Metabolic Disease, 43(4), 694–700. 10.1002/jimd.12237

Parrado, A., Rubio, G., Serrano, M., De la Morena-Barrio, M. E., Ibáñez-Micó, S., Ruiz-Lafuente, N., Schwartz-Albiez, R., Esteve-Solé, A., Alsina, L., Corral, J., & Hernández-Caselles, T. (2022). Dissecting the transcriptional program of phosphomannomutase 2-deficient cells: Lymphoblastoide B cell lines as a valuable model for congenital disorders of glycosylation studies. Glycobiology, 32(2), 84–100. 10.1093/glycob/cwab087

Radenkovic, S., Budhraja, R., Klein-Gunnewiek, T., King, A. T., Bhatia, T. N., Ligezka, A. N., Driesen, K., Shah, R., Ghesquière, B., Pandey, A., Kasri, N. N., Sloan, S. A., Morava, E., & Kozicz, T. (2024). Neural and metabolic dysregulation in PMM2-deficient human in vitro neural models. Cell Reports, 43(3), 113883. 10.1016/j.celrep.2024.113883

Radenkovic, S., Ligezka, A. N., Mokashi, S. S., Driesen, K., Dukes-Rimsky, L., Preston, G., Owuocha, L. F., Sabbagh, L., Mousa, J., Lam, C., Edmondson, A., Larson, A., Schultz, M., Vermeersch, P., Cassiman, D., Witters, P., Beamer, L. J., Kozicz, T., Flanagan-Steet, H., … Morava, E. (2023). Tracer metabolomics reveals the role of aldose reductase in glycosylation. Cell Reports Medicine, 4(6). 10.1016/j.xcrm.2023.101056

Radenkovic, S., Preston, G., Budhraja, R., Muffels, I., Ligezka, A. N., Hrstka, R., Staff, N. P., Balakrishan, B., Shah, R., Verberkmoes, S., Shammas, I., Bosnyak, I., Stiers, K. M., Lai, K., Beamer, L. J., Pandey, A., Morava, E., & Kozicz, T. (2025). PGM1 deficiency disrupts sarcomere and mitochondrial function in a stem-cell cardiomyocyte model. 10.1101/2025.07.01.662580

Stillwell, W., Jenski, L. J., Thomas Crump, F., & Ehringer, W. (1997). Effect of docosahexaenoic acid on mouse mitochondrial membrane properties. Lipids, 32(5), 497–506. 10.1007/s11745-997-0064-6

Sun, J., Zhang, Y., Yu, W., Fu, H., Lin, N., Yu, F., Chen, X., Mao, J., & Hu, L. (2025). Cysteine variants in PMM2 lead to protein instability and higher sensitivity to oxidative stress in PMM2-CDG. International Journal of Biological Macromolecules, 305, 140865. 10.1016/j.ijbiomac.2025.140865

Verheijen, J., Tahata, S., Kozicz, T., Witters, P., & Morava, E. (2020). Therapeutic approaches in Congenital Disorders of Glycosylation (CDG) involving N-linked glycosylation: an update. Genetics in Medicine, 22(2), 268–279. 10.1038/s41436-019-0647-2

Wessels, H. J. C. T., Kulkarni, P., van Dael, M., Suppers, A., Willems, E., Zijlstra, F., Kragt, E., Gloerich, J., Schmit, P.-O., Pengelley, S., Marx, K., van Gool, A. J., & Lefeber, D. J. (2024). Plasma glycoproteomics delivers high-specificity disease biomarkers by detecting site-specific glycosylation abnormalities. Journal of Advanced Research, 61, 179–192. 10.1016/j.jare.2023.09.002

Wishart, D. S., Knox, C., Guo, A. C., Shrivastava, S., Hassanali, M., Stothard, P., Chang, Z., & Woolsey, J. (2006). DrugBank: a comprehensive resource for in silico drug discovery and exploration. Nucleic Acids Research, 34(Database issue), D668–72. 10.1093/nar/gkj067

Zdrazilova, L., Rakosnikova, T., Himmelreich, N., Ondruskova, N., Pasak, M., Vanisova, M., Volfova, N., Honzik, T., Thiel, C., & Hansikova, H. (2023). Metabolic adaptation of human skin fibroblasts to ER stress caused by glycosylation defect in PMM2-CDG. Molecular Genetics and Metabolism, 139(4), 107629. 10.1016/j.ymgme.2023.107629

Zhou, X., Wang, M., Katsyv, I., Irie, H., & Zhang, B. (2018). EMUDRA: Ensemble of Multiple Drug Repositioning Approaches to improve prediction accuracy. Bioinformatics, 34(18), 3151–3159. 10.1093/bioinformatics/bty325

